# Structural basis for Ca_V_α_2_δ:gabapentin binding

**DOI:** 10.1101/2022.12.05.519047

**Authors:** Zhou Chen, Abhisek Mondal, Daniel L. Minor

## Abstract

Gabapentinoid drugs for pain and anxiety act on the Ca_V_α_2_δ-1 and Ca_V_α_2_δ-2 subunits of high-voltage activated calcium channels (Ca_V_1s and Ca_V_2s). Here, we present the cryo-EM structure of the gabapentin-bound brain and cardiac Ca_V_1.2/Ca_V_β_3_/Ca_V_α_2_δ-1 channel. The data reveal a binding pocket in the Ca_V_α_2_δ-1 dCache1 domain that completely encapsulates gabapentin and define Ca_V_α_2_δ isoform sequence variations that explain gabapentin binding selectivity of Ca_V_α_2_δ-1 and Ca_V_α_2_δ-2.

---

Gabapentin (GBP) (1-(aminomethyl)cyclohexane acetic acid; Neurontin)^1^ and related gabapentinoid drugs pregabalin (Lyrica)^2,3^ and mirogabalin (Tarlige)^4,5^, are widely used to treat post-herpetic neuralgia, diabetic neuropathy, fibromyalgia, epilepsy, restless leg syndrome, and generalized anxiety disorder. These drugs bind to high-voltage activated calcium channel (Ca_V_) Ca_V_α_2_δ-1 and Ca_V_α_2_δ-2 subunits^3,4,6-8^, but not to the related Ca_V_α_2_δ-3 and Ca_V_α_2_δ-4 isoforms^2,9,10^. Gabapentinoid binding to Ca_V_α_2_δ-1 and Ca_V_α_2_δ-2 is thought to affect neuronal excitability by impairing Ca_V_ surface membrane expression^3,7,8^ through a mechanism involving Rab11a endosomal recycling^11,12^. Although the GBP binding site has been identified^13^, structural details of Ca_V_α_2_δ:GBP interactions have not yet been defined.

High voltage-gated calcium channels (Ca_V_1 and Ca_V_2)^14,15^ are multi-subunit voltage-gated ion channels comprising three key components, the pore-forming Ca_V_α^114,15^, cytoplasmic Ca_V_β^16^, and extracellular Ca_V_α_2_δ^1,17^ subunits. Recent, cryo-EM structural studies of the Ca_V_1.2/Ca_V_β_3_/Ca_v_α_2_δ-1 channel complex^13^ revealed a Ca_V_α_2_δ-1-bound L-leucine, a known Ca_V_α_2_δ ligand^1,18,19^ and gabapentin competitor^1,20^, that identified the gabapentinoid binding site^1,17^. Here, we present the 3.1Å cryo-EM structure of the Ca_V_1.2/Ca_V_β_3_/Ca_v_α_2_δ-1 channel bound to GBP. The data show that, similar to L-Leu^13^, GBP occupies the L-Leu binding site in the first subdomain of the Ca_v_α_2_δ-1 dCache1 (<u>Ca</u>^2+^ channel and <u>che</u>motaxis receptor) domain^21^. Contact analysis identifies five binding site residues that differ between the GBP-sensitive isoforms, Ca_V_α_2_δ- 1 and Ca_V_α_2_δ-2^3,4,6-8^, and the GBP-insensitive isoforms, Ca_V_α_2_δ-3 and Ca_V_α_2_δ-4^2,9,10^. These yield steric clashes with GBP and cause binding site polarity changes that rationalize the GBP-insensitivity of Ca_V_α_2_δ-3 and Ca_V_α_2_δ-4 isoforms.

Structure determination of a recombinant Ca_V_1.2/Ca_V_β_3_/Ca_v_α_2_δ-1 channel complex^13^ comprising human Ca_V_1.2(βC) (186 kDa), rabbit Ca_V_β_3_ (54 kDa), and rabbit Ca_v_α_2_δ-1 (125 kDa) in the presence of 11.7 mM GBP revealed a tripartite channel assembly (∼370 kDa) (Fig.1a) at an overall resolution of 3.1Å (Fig. S1, S2a-c, Table S1) largely similar to the L-Leu bound structure^13^ (RMSD_Cα_ = 0.749Å). As previously observed^13^, the sample also contained a chaperone:channel complex of the ER membrane protein complex (EMC)^22-24^, Ca_V_1.2(βC), and Ca_V_β_3_ (Fig. S1, S2d-e). The overall structure of the Ca_V_1.2/Ca_V_β_3_/Ca_v_α_2_δ-1:GBP complex is similar to other Ca_V_1 and Ca_V_2 structures^13,25-27^. Ca_v_α_2_δ-1 has a multidomain architecture built from two double Cache domains^28^, dCache1 and dCache2, and a von Willebrand factor type A (VWA) domain^25,28,29^ (Figs. 1a and S3a). Importantly, the excellent Ca_v_α_2_δ-1 local resolution (2.0-2.5Å) and map quality (Figs. 1b-d, S2b-c, S3a-f) allowed detailed comparison of the dCache1 domain with the L-Leucine bound structure^13^. Map comparison (Fig. 1b-d) showed clear differences in the binding pocket density, including a shape not present in the L-Leu bound maps (Fig. 1b-d. that had features that could be attributed to the GBP cyclohexyl ring defining the GBP binding site.

**Figure 1.**
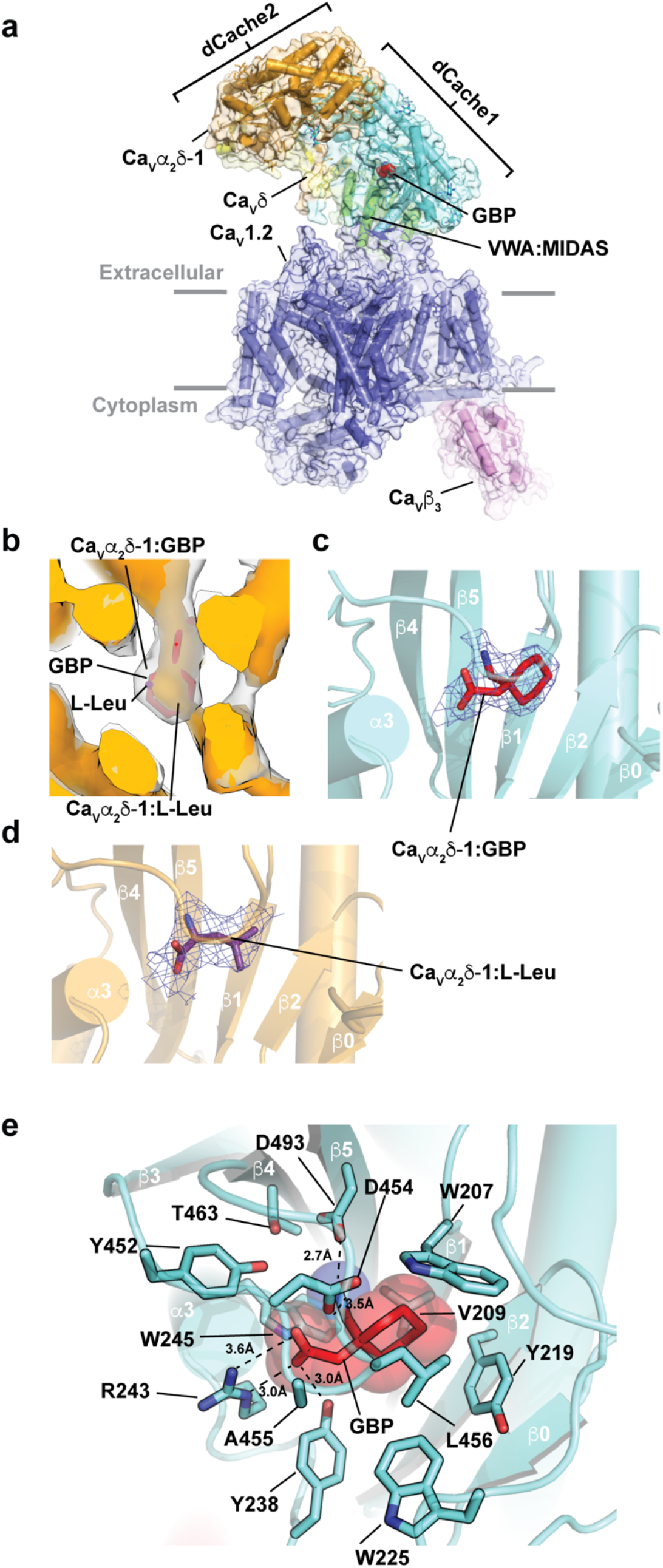
Structure of the Ca_V_1.2(ΔC)/Ca_V_β_3_/Ca_v_α_2_δ-1:GBP complex. **a**, Side view of the Ca_V_1.2(ΔC)/Ca_V_β_3_/Ca_V_α_2_δ-1:GBP complex. Subunits are colored: Ca_V_1.2 (slate) and Ca_V_β_3_ (violet). Ca_V_α_2_δ domains are colored as: dCache1(aquamarine), dCache2 (orange), VWA:MIDAS (green), and Ca_V_δ (yellow). GBP (red) is shown as space filling. Grey bars denote the membrane. **b**, Comparison of Ca_V_α_2_δ-1 dCache1 binding sites. Map comparison of the dCache1 ligand binding site for the GBP complex (clear) and L-Leu complex^13^ (orange). GBP is shown as red sticks. **c**, and **d**, Ligand density for **c**, GBP (10σ) (red), and **d**, L-Leu (7σ) (violet purple) in dCache1 domains are shown as cartoons (aquamarine and orange, respectively). **e**, Ca_V_α_2_δ-1 dCache1 ligand binding site details. GBP (red) and contacting Ca_V_α_2_δ−1 sidechains (greencyan) are shown as sticks.

Similar to L-Leu, GBP occupies a pocket in the first Ca_v_α_2_δ-1 dCache1 β-barrel lined by Trp207, Val209, Tyr219, Trp225, Tyr238, Arg243, Trp245, Tyr452, Asp454, Ala455, Leu456, Thr463, and Asp493 (Figs. 1e and S3f) that is closed to solvent access. In line with the similar affinities of the two ligands^18^, there are no large conformational differences in the dCache1 binding site relative to the Ca_v_α_2_δ-1:L-Leu complex (RMSD_Cα_ = 0.155Å) (Fig. S4a). The structure shows that GBP buries a larger total surface area than L-Leu (409Å^2^ vs. 367Å^2^ for GBP and L-Leu, respectively) and that the different sizes and shapes of the ligands alter the details of the hydrogen bond network surrounding the carboxylate and amino groups found in both ligands (Fig. S4b-c). As with the L-Leu complex, Arg243 (Arg217 in some numbering schemes), a key residue for GBP binding to Ca_v_α_2_δ-1^30^, is central to the coordination of the GBP carboxylate and makes bidentate interactions to the ligand through its guanidinium group (Figs. 1c, S4b). The importance of this interaction is supported the observation that an R→A change at this site is known to eliminate GBP binding^3,30^, abolish the analgesic effects of GBP and mirogabalin on pain^3,31^, and mitigate the effects of pregabalin on seizure and anxiety^32,33^. The Asp493 sidechain makes a salt bridge with the GBP amino group that is shorter than the similar interaction in the L-Leu complex (2.7Å vs. 3.2Å, GBP and Leu, respectively) (Fig. S4b-c). Disruption of this interaction by mutation to alanine strongly reduces the effect of GBP on Ca_V_ function^28^. Together with the previously demonstrated potent effects of Arg243^3,30-33^ and Asp493^28^ alanine mutations on gabapentinoid binding and function, the structure underscores the importance of the dCache1 hydrogen bond network that coordinates the amino and carboxylate moieties shared by amino acid and gabapentinoid Ca_v_α_2_δ ligands.

The first subdomain of dCache1 repeats is a common binding site for free amino acids in many archaeal, bacterial, and eukaryotic dCache domain containing proteins^28,34^, as exemplified by structures of the *Psuedomonas aeruginosa* chemoreceptor PctA^34^. The overall fold of the first Ca_v_α_2_δ-1 dCache1 domain is largely similar to the amino acid binding dCache1 domain of bacterial PctA^34^ (RMSD_Cα_ = 4.123Å). However, structural comparison reveals key differences including the long Ca_v_α_2_δ-1 β2-α3 loop, absent from bacterial dCache domains (Fig. S4d), the relative position of the β3-β4 loop that covers the ligand binding pocket, and positional changes in the residues that encircle the ligand (Fig. S4e). Nevertheless, Ca_v_α_2_δ-1 uses the signature dCache1 domain amino acids that recognize the carboxylate (YxxxxRxW) and amino (Y[x_∼27-34_]D) groups of various amino acid derived ligands in bacterial dCache domains^28^ to coordinate the GBP and L-Leu carboxylate and amino and carboxylate moieties (Y238, R243, W245 and Y452, D493), respectively, through common hydrogen bond networks (Fig. S4b-c). This network is augmented in Ca_v_α_2_δ-1 by W225, a β2-α3 loop residue (Fig. 2a) that contributes to carboxylate coordination (Fig. S4b-c). In bacterial dCache domains^34^ the equivalent of the β3-β4 loop that covers the GBP binding site moves to open access to the amino acid binding pocket, suggesting that similar Ca_v_α_2_δ-1 dCache1 motions could provide a means for L-Leu, GBP, and other gabapentinoids to access the Ca_v_α_2_δ-1 binding pocket.

**Figure 2.**
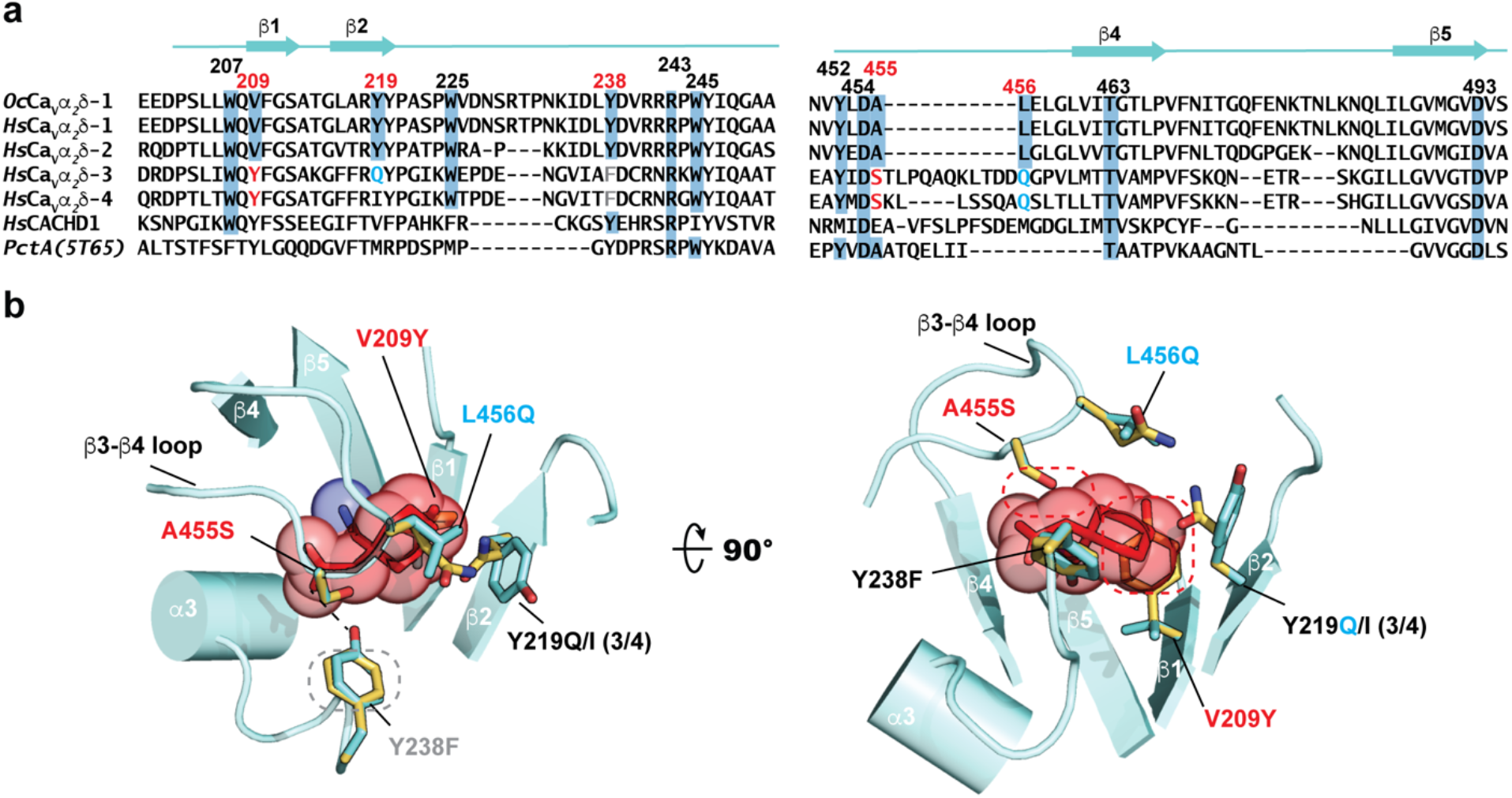
Comparison of Ca_V_α_2_δ-1 GBP binding sites. **a**, Sequence comparison of rabbit *Oc*Ca_V_α_2_δ-1 (NCBI: NP_001075745.1), human Ca_V_α_2_δ-1 (NCBI: NP_001353796.1), Ca_V_α_2_δ-2 (NCBI: NP_006021.2), Ca_V_α_2_δ-3 (NCBI: NP_060868.2), and Ca_V_α_2_δ-4 (NCBI: NP_758952.4), CACHD1 (NCBI: NP_065976.3), and bacterial PctA (PDB:5T65)^34^ dCache1 domain sequences. Numbers indicate residues interacting with L-Leu and GBP. Red numbers indicate sites that differ between GBP-sensitive and GBP-insensitive isoforms. Residue colors indicate steric clash (red), hydrogen bond loss (grey), and polarity changes (blue) between GBP-sensitive and GBP-insensitive isoforms. **b**, Structural context for amino acid differences between GBP-sensitive (aquamarine) and GBP-insensitive (yellow) Ca_V_α_2_δ-1 Cache1 ligand binding sites. GBP-insensitive residues are modeled on the Ca_V_α_2_δ-1:GBP structure. Dashed ovals denote hydrogen bond loss (grey) and steric clashes (red). (3/4) indicates amino acid changes for Ca_V_α_2_δ-3 and Ca_V_α_2_δ-4.

The four different mammalian Ca_v_α_2_δ isoforms share a common structure^28^, yet gabapentin and related gabapentinoids bind to Ca_V_α_2_δ-1 and Ca_V_α_2_δ-2^3,4,6-8^ but not Ca_V_α_2_δ-3 and Ca_V_α_2_δ-4^2,9,10^. Structure-based sequence comparisons identify five sites in the GBP binding pocket that differ between the gabapentinoid-sensitive (Ca_V_α_2_δ-1 and Ca_V_α_2_δ-2^3,4,6-8^) and gabapentinoid-insensitive (Ca_V_α_2_δ-3 and Ca_V_α_2_δ-4^2,9,10^) Ca_V_α_2_δ isoforms (Fig. 2a). Mapping these on the Ca_v_α_2_δ-1:GBP structure (Fig. 2b) reveals numerous alterations expected to interfere with GBP binding to Ca_V_α_2_δ-3 and Ca_V_α_2_δ-4 including: 1) loss of a hydrogen bond donor to the GBP carboxylate (Y238F), 2) introduction of steric clashes, (V209Y and A455S), one of which (V209Y) would occupy the same space as the GBP cyclohexyl ring, and 3) changes in contact residue size (Y219I) and polarity (Y219Q, L456Q, A455S) that reshape binding pocket physiochemical characteristics. The Ca_V_α_2_δ related protein CACHD1 also lacks most of the key GBP binding residues (Fig. 2a) indicating that the effects of this protein on Ca_V_ surface expression^35^ are likely to be GBP-independent. Taken together, the multiple changes between the GBP-sensitive and GBP-insensitive isoforms that remove hydrogen bonds, introduce steric clashes, and reduce the hydrophobicity of the binding pocket provide a structural rationalization for the differences in gabapentinoid binding properties among Ca_V_α_2_δ isoforms^2-4,7-10^.

Binding of GBP and gabapentinoids to the Ca_V_α_2_δ subunit of Ca_V_s is critical for the pharmacological effects of this drug class^3,31-33^. Structural definition of the Ca_V_α_2_δ gabapentinoid binding site provides a platform to dissect the mechanisms by which these drugs affect Ca_V_ function and should guide the development of next generation Ca_V_α_2_δ-directed drugs for treatment of pain and anxiety.

## Supporting information

Supplementary Figs. S1-S4, Table S1, Movie legend M1, and References

Movie S1 CaVa2d-1 ligand binding site cryo-EM density comparison.

## Acknowledgements

We thank D. Bulkley for technical help and K. Brejc for comments on the manuscript. This work was supported by grant NIH R01 HL080050 to D.L.M.

## Author Contributions

Z.C., A.M., and D.L.M. conceived the study and designed the experiments. Z.C. expressed and characterized the samples. Z.C. and A.M., collected and analyzed cryo-EM data. Z.C. and A.M. built and refined the atomic models. D.L.M. analyzed data and provided guidance and support. Z.C., A.M., and D.L.M. wrote the paper.

## Data and materials availability

Ca_v_1.2(ΔC)/Ca_v_β_3_/Ca_v_α_2_δ-1:GBP coordinates and maps (PDB:8FD7; EMD-29004) and map of the EMC:Ca_v_1.2(ΔC)/Ca_v_β_3_ complex (EMD-29006) are deposited with the RCSB and will be released upon publication.

Requests for material to D.L.M.

## Materials and Methods

### Expression and purification of human Ca_v_1.2

Expression and purification of Ca_V_1.2(ΔC)/Ca_V_β_3_/Ca_V_α_2_δ-1 was done as previously described ^13^ in HEK293S GnTI^-^ (ATCC) ‘Ca_V_β_3_-stable’ expressing rabbit Ca_V_β_3_ (477 residues, Uniprot P54286) bearing a Strep*-*tag II sequence^36^ and codon-optimized cDNAs of human Ca_v_1.2 bearing a C-terminal truncation at residue 1648 (denoted Ca_V_1.2(ΔC)), a site 13 residues after the end of the IQ domain (Δ1649-2138, Uniprot Q13936-20, 1,648 residues) followed by a 3C protease cleavage site, monomeric enhanced Green Fluorescent Protein (mEGFP), and a His_8_ tag, rabbit Ca_v_α_2_δ-1 (1,105 residues, Uniprot P13806-1), modified pFastBac expression vectors having the polyhedrin promoter replaced by a mammalian cell active CMV promoter^37^. All constructs were sequenced completely.

Chemically competent DH10EmBacY (Geneva Biotech) were used to generate the recombinant bacmid DNA, which was then used to transfect *Spodoptera frugiperda* (Sf9) cells to make baculoviruses for each subunit^38^. Ca_v_1.2 was expressed in Ca_V_β_3_-stable cells together with Ca_v_α_2_δ-1 using a baculovirus transduction-based system^38^. Ca_V_β_3_-stable cells were grown in suspension at 37°C supplied with 8% CO_2_ in FreeStyle 293 Expression Medium (Gibco) supplemented with 2% fetal bovine serum (FBS, Peak Serum), and were transduced with 5% (v/v) baculovirus for each target subunit when cell density reached ∼2.5 × 10^6^ cells per ml. 10 mM sodium butyrate was added to cell culture 16-24 h post-transduction and the cells were subsequently grown at 30°C. Cells were harvested 48 h post-transduction by centrifugation at 5,000*g* for 30 min. The pellet was washed with Dulbecco’s phosphate buffered saline (DPBS) (Gibco) and stored at -80°C.

A cell pellet (from ∼3.6 L culture) was resuspended in 200 ml of resuspension buffer containing 0.3 M sucrose, 1 mM ethylenediaminetetraacetic acid (EDTA), 10 mM Tris-HCl, pH 8.0 supplemented with 1 mM phenylmethylsulfonyl fluoride (PMSF) and 4 Pierce protease inhibitor tablets (Thermo scientific), then stirred gently on a Variomag magnetic stirrer MONO DIRECT (Thermo Scientific) at 4°C for 30 min. The membrane fraction was collected by centrifugation at 185,500*g* for 1 h and subsequently solubilized in 200 ml of solubilization buffer (buffer S) containing 500 mM NaCl, 5% glycerol (v/v), 0.5 mM CaCl2, 20 mM Tris-HCl, pH 8.0, and supplemented with 1% (w/v) glycol-diosgenin (GDN) and rotated on an Orbitron rotator II (speed mode S) (Boekel Scientific) at 4°C for 2 h. The supernatant, collected by centrifugation at 185,500*g* for 1 h, was diluted with an equal volume of buffer S to a final concentration of 0.5% GDN and incubated with anti-GFP nanobody Sepharose resin^39^ at 4°C overnight. The resin was loaded on an Econo-Column chromatography column (BioRad) and was then washed stepwise with 20 column volumes (CV) of buffer S supplemented with 0.1% (w/v) GDN, 20 CV of buffer S supplemented with 0.02% (w/v) GDN, and 20 CV of elution buffer (buffer E) containing 150 mM NaCl, and 0.5 mM CaCl_2_, 0.02% (w/v) GDN 20 mM Tris-HCl pH 8.0. The protein was eluted with 3C protease^40^ and subsequently incubated at 4°C for 2 h with 4 mL of *Strep*-tactin Superflow Plus beads (Qiagen) pre-equilibrated with buffer E. The beads were washed with 20 column volumes of buffer E and the protein was eluted with buffer E supplemented with 2.5 mM desthiobiotin. The eluent was concentrated using an Amicon Ultra-15 100-kDa cut-off centrifugal filter unit (Merck Millipore) before purification on a Superose 6 Increase 10/300 GL gel filtration column (GE Healthcare) pre-equilibrated in buffer E. Concentrated Ca_v_1.2(ΔC)/Ca_v_β_3_/Ca_v_α_2_δ-1 sample was immediately incubated with gabapentin (final concentration 2 mg ml^-1^, 11.7 mM) (Sigma-Aldrich) on ice for four hours prior to cryo-EM sample preparation, denoted Ca_v_1.2(ΔC)/Ca_v_β_3_/Ca_v_α_2_δ-1:GBP.

### Sample preparation and cryo-EM data acquisition

For cryo-EM, 3.5 μl of 2.0 mg ml^-1^ Ca_v_1.2(ΔC)/Ca_v_β_3_/Ca_v_α_2_δ-1:GBP was applied to Quantifoil R1.2/1.3 300 mesh Au holey-carbon grids, blotted for 4-6 s at 4°C and 100% humidity using a FEI Vitrobot Mark IV (Thermo Fisher Scientific), and plunge frozen in liquid ethane. Cryo-EM grids were screened on a FEI Talos Arctica cryo-TEM (Thermo Fisher Scientific) (at University of California, San Francisco (UCSF) EM Facility) operated at 200 kV and equipped with a K3 direct detector camera (Gatan), and then imaged on a 300 kV FEI Titan Krios microscope (Thermo Fisher Scientific) with a K3 direct detector camera (Gatan) (UCSF). Cryo-EM datasets were collected in super-resolution counting mode mode at a nominal magnification of 105,000x with a super-resolution pixel size of 0.4175 Å (physical pixel size of 0.835 Å) using SerialEM^41^. Images were recorded with a 2.024 s exposure over 81 frames with a dose rate of 0.57 e^−^ Å^−2^ per frame. The defocus range was set from -0.9 μm to -1.7 μm.

### Imaging processing and 3D reconstruction

A total of 26,928 movies were collected for Ca_v_1.2(ΔC)/Ca_v_β_3_/Ca_v_α_2_δ-1:GBP. Initial image processing was carried out in cryoSPARC-3.3^42^. Raw movies were corrected for motion and Fourier binned by a factor of two (final pixel size of 0.834 Å) with the patch motion job. Contrast transfer function (CTF) parameters of the resulting micrographs were estimated with the Patch CTF estimation program in cryoSPARC-3.3. Particles were picked by blob picking, extracted using a box size of 440 pixel (2x binned to 220 pixel), and 2D-classified using a mask diameter of 260 Å. Selected particles represented by good 2D classes were subjected to one round of *ab initio* reconstruction and heterogeneous refinement with C1 symmetry. Particles having reasonable 3D reconstructions (as judged by the Fourier shell correlation (FSC) curve) were re-extracted to physical pixel size and subjected to iterative rounds of *ab initio* reconstruction and heterogeneous refinement. Further, Non-uniform refinements were performed to achieve high-resolution reconstruction.

To improve the Ca_v_α_2_δ-1 3D reconstruction, multibody refinement was carried out in RELION-3.1^43^. In total, 259,107 refined particles for the Ca_V_1.2(ΔC)/Ca_V_β_3_/Ca_V_α_2_δ-1:GBP complex were exported from cryoSPARC-3.3 to RELION-3.1 using the csparc2star.py (UCSF pyem v0.5. Zenodo) suite of conversion scripts (https://doi.org/10.5281/zenodo.3576630). Following a 3D refinement in RELION-3.1 using the refined map from cryoSPARC-3.3 and the exported particles, an overall 3.1 Å EM density map (consensus map) was obtained (see Supplementary Fig. 1 for processing flow charts). Multibody refinement was performed in RELION-3.1 to improve the features of the lumenal domain and the transmembrane region for the Ca_V_1.2(ΔC)/Ca_V_β_3_/Ca_V_α_2_δ-1:GBP complex. phenix.combine_focused_maps program was used to combine the improved features of the segments from multibody refinement and those of the consensus map to obtain the final map with best features for the Ca_V_1.2(ΔC)/Ca_V_β_3_/Ca_V_α_2_δ-1:GBP complex^44^.

As with the prior report^13^, purification of Ca_V_1.2(ΔC)/Ca_V_β_3_/Ca_V_α_2_δ-1 yielded a sample that also had a substantial portions of particles (383,185 refined particles) comprising the EMC: Ca_V_1.2(ΔC)/Ca_V_β_3_ complex. These refined particles were exported from cryoSPARC-3.3 to RELION-3.1 and subsequent 3D refinement resulted in a 3.1 Å consensus map. Comparison of this 3.1 Å consensus map for the EMC: Ca_V_1.2(ΔC)/Ca_V_β_3_ complex with the one reported from the previous study^13^ revealed no apparent conformational difference (cross correlation = 0.9554), thus, model was not docked for this complex.

### Model building and refinement

phenix.dock_in_map ^44^ was used to dock the Ca_v_1.2(ΔC)/Ca_v_β_3_/Ca_v_α_2_δ-1 model (PDB:8EOG) into the Ca_v_1.2(ΔC)/Ca_v_β_3_/Ca_v_α_2_δ-1:GBP map. Docked model and maps were manually checked and fitted in COOT^45^. Iterative structure refinement and model building were performed using phenix.real_space_refine program^44^. Restraint files necessary for refinement were generated using phenix.elbow^44,46^. Final statistics of 3D reconstruction and model refinement can be found in (Table S1). The per-residue B-factors, after final refinement against the overall map, were rendered on the refined model and presented in (Fig. S2c). The final models were evaluated using MolProbity^47^. All figures and movies were generated using ChimeraX^48^ and Pymol package (http://www.pymol.org/pymol). Close-contact interaction analysis were performed using LIGPLOT and DIMPLOT^49,50^.

## Supplementary material

Supplementary material contains Figures S1-S4, Movie S1, Table S1, and references.

