## Supplementary Figs. S1-S4, Table S1, Movie legend M1, and References for "Structural basis for Ca_V_α_2_δ:gabapentin binding"

4 December 2022

**Structural basis for $Ca_V\alpha_2\delta$:gabapentin binding**

[5]Molecular Biophysics and Integrated Bio-imaging Division
Lawrence Berkeley National Laboratory, Berkeley, CA 94720 USA

[§]Equal contributions

**Keywords:** voltage-gated calcium channel, $Ca_V\alpha_2\delta$, drug binding, gabapentin

**Figure S1** **Chen, Mondal & Minor**

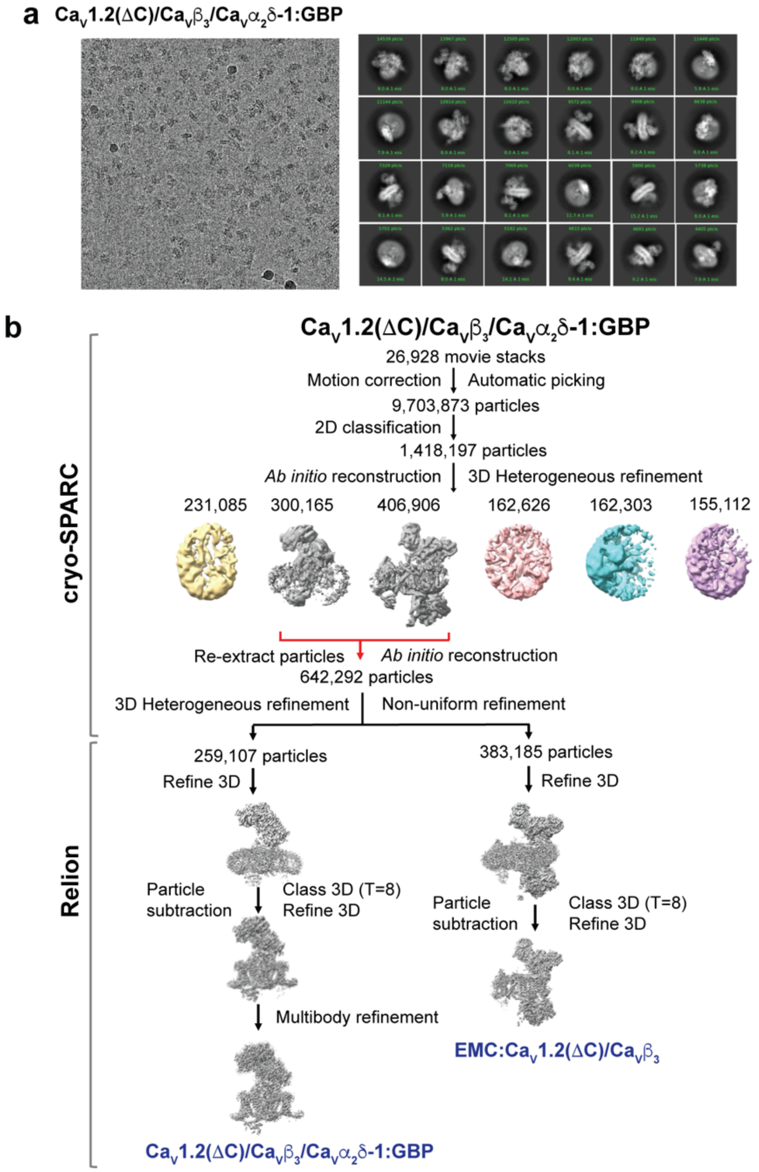


**Figure S1 $Ca_V1.2(\Delta C)/Ca_V\beta_3/Ca_V\alpha_2\delta$-1:GBP Cryo-EM analysis a,** Exemplar $Ca_V1.2(\Delta C)/Ca_V\beta_3/Ca_V\alpha_2\delta$-1:GBP electron micrograph (~105,000x magnification), and 2D class averages. **b**, Workflow for electron microscopy data processing for $Ca_V1.2(\Delta C)/Ca_V\beta_3/Ca_V\alpha_2\delta$-1:GBP sample. Initial cryoSPARC-3.2 *Ab initio* reconstruction identified a population of particles containing the $Ca_V1.2(\Delta C)/Ca_V\beta_3/Ca_V\alpha_2\delta$-1 and EMC:$Ca_V1.2(\Delta C)/Ca_V\beta_3$ complexes, similar to prior studies[1]. Red arrows indicate the two classes that were re-extracted, subjected to multiple rounds of 3D heterogeneous classification, and exported from cryoSPARC-3.2 for further 3D refinement in RELION-3.1. Particle subtraction was performed for both the refined maps in Relion-3.1 followed by 3D classification with single class and 3D refinement to get the final consensus maps. Multibody refinement was performed to enhance features of $Ca_V\alpha_2\delta$-1, which was used for the $Ca_V1.2(\Delta C)/Ca_V\beta_3/Ca_V\alpha_2\delta$-1:GBP composite map. The composite map was used for model building and refinement.

**Figure S2** **Chen, Mondal & Minor**

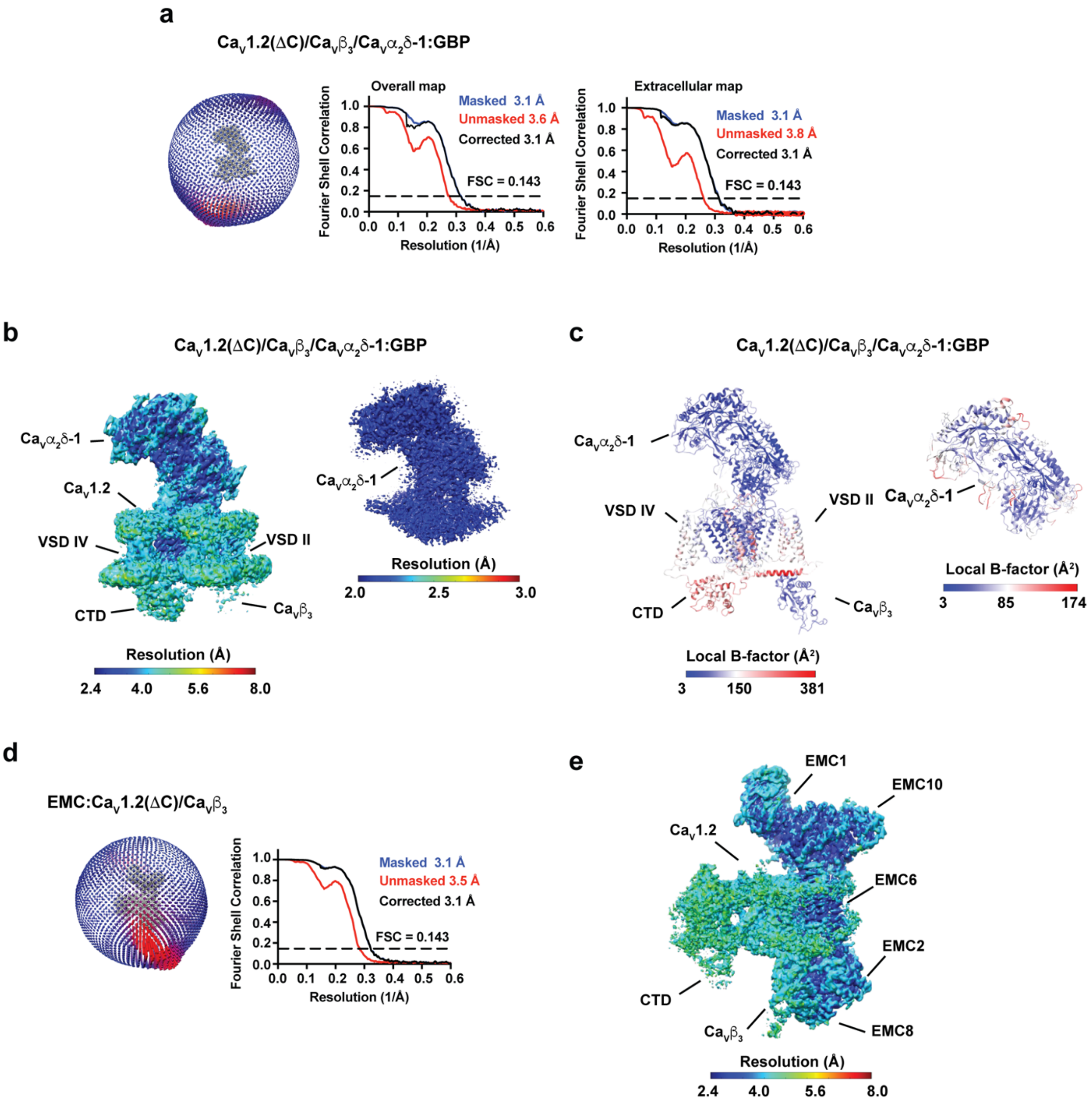


**Figure S2 $Ca_V1.2(\Delta C)/Ca_V\ \beta_3/Ca_V\alpha_2\delta$-1:GBP map and model quality a,** Particle distribution plot and gold-standard Fourier shell correlation (FSC) curve for the overall $Ca_V1.2(\Delta C)/Ca_V\ \beta_3/Ca_V\alpha_2\delta$-1:GBP complex map and the extracellular map containing $Ca_V\alpha_2\delta$-1:GBP. **b,** local resolution for the overall $Ca_V1.2(\Delta C)/Ca_V\ \beta_3/Ca_V\alpha_2\delta$-1:GBP map and the extracellular map containing $Ca_V\alpha_2\delta$-1:GBP. **c,** local B-factor for the overall $Ca_V1.2(\Delta C)/Ca_V\ \beta_3/Ca_V\alpha_2\delta$-1:GBP model and the $Ca_V\alpha_2\delta$-1:GBP subunit. **d,** Particle distribution plot and gold-standard Fourier shell correlation (FSC) curve for the EMC:$Ca_V1.2(\Delta C)/Ca_V\beta_3$

complex from the $Ca_V1.2(\Delta C)/Ca_V\ \beta_3/Ca_V\alpha_2\delta$-1:GBP sample. **e,** EMC:$Ca_V1.2(\Delta C)/Ca_V\beta_3$ complex local resolution. Select elements of each complex are labeled.

**Figure S3** **Chen, Mondal & Minor**

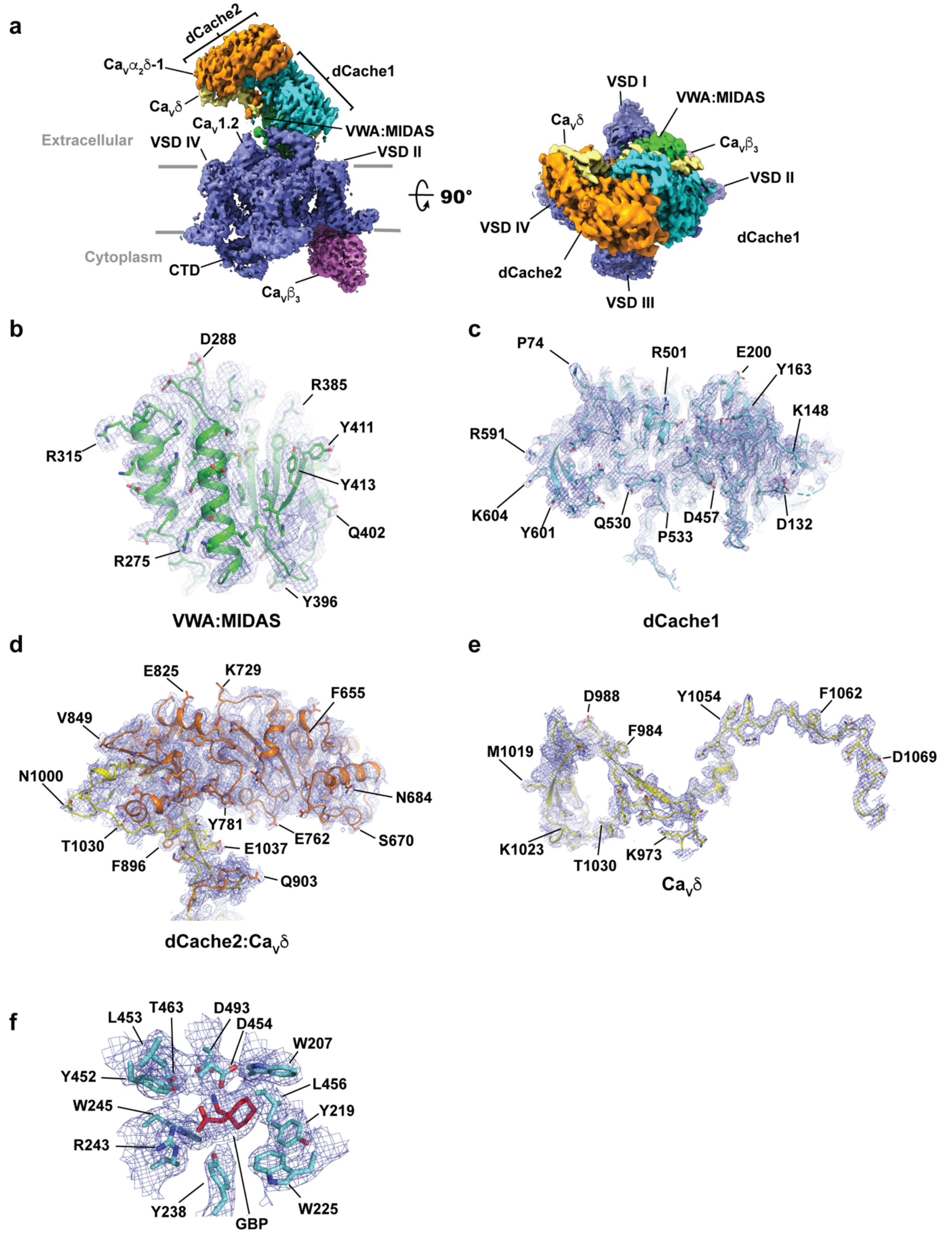


**Figure S3 Ca$_V$1.2(ΔC)/Ca$_V$β$_3$/Ca$_V$α$_2$δ-1:GBP Cryo-EM maps. a,** Ca$_V$1.2(ΔC)/Ca$_V$β$_3$/Ca$_V$α$_2$δ-1 side view (left) and extracellular (right) views. Subunits are colored: Ca$_V$1.2 (slate) and Ca$_V$β$_3$

(violet). $Ca_V\alpha_2\delta$ domains are colored as: dCache1 (aquamarine), dCache2 (orange), VWA:MIDAS (green), and $Ca_V\delta$ (yellow). Grey bars denote the membrane. **b-e,** $Ca_V\alpha_2\delta$-1 subdomain representative maps for **b,** VWA:MIDAS domain (green), **c,** dCache1, **d,** dCache2:$Ca_V\delta$. Part of $Ca_V\delta$ completes the second $\beta$-barrel subdomain of dCache2, **e,** $Ca_V\delta$. **f,** GBP binding site. Maps are rendered at 9-10$\sigma$**.**

**Figure S4** **Chen, Mondal & Minor**

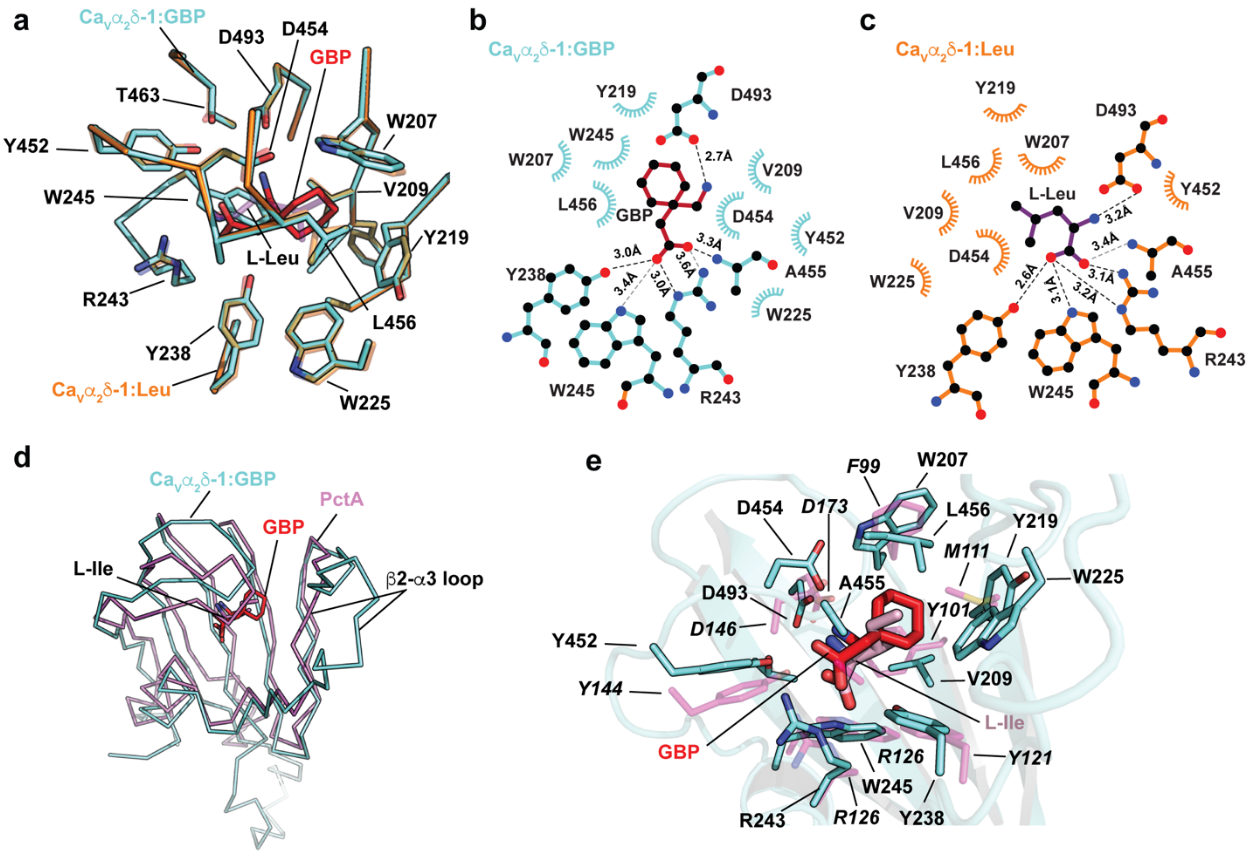


**Figure S4 $Ca_V\alpha_2\delta$-1 GBP binding site analysis and comparisons. a,** Superposition of the $Ca_V\alpha_2\delta$-1:GBP (aquamarine) and $Ca_V\alpha_2\delta$-1:L-Leu (orange) (PDB:8EOG)[1] binding sites. GBP is red. L-Leu is purple. **b** and **c,** LigPLOT [2] diagrams of the **b,** $Ca_V\alpha_2\delta$-1:GBP (aquamarine) and **c,** $Ca_V\alpha_2\delta$-1:L-Leu (orange) (PDB:8EOG)[1] binding sites showing hydrogen bonds and ionic interactions (dashed lines) and van der Waals contacts ≤ 5Å. GBP is red. L-Leu is purple. **d,** Superposition of the first dCache1 repeats from $Ca_V\alpha_2\delta$-1:GBP (aquamarine) and the PctA:L-Ile complex (magenta) (PDB: 5T65)[3]. GBP is red. **e,** Closeup view of superposition from 'd' showing ligand contact residues. $Ca_V\alpha_2\delta$-1 is shown as a cartoon. GBP is red. Corresponding sidechains of PctA are magenta. L-Ile form the PctA complex is pink. PctA residues are labeled in italics.

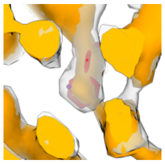

**Movie S1 $Ca_V\alpha_2\delta$-1 ligand binding site cryo-EM density comparison.** Movie shows superposition of maps for the $Ca_V\alpha_2\delta$-1:GBP (13.9$\sigma$) (clear) and $Ca_V\alpha_2\delta$-1:L-Leu (7.5$\sigma$)(orange) (EMD-28375)[1] GBP (red) and L-Leu (purple) are shown as sticks.

**Table S1 Statistics for data collection, refinement, and validation**

| | $Ca_V1.2(\Delta C)/Ca_V\beta_3/Ca_V\alpha_2\delta$-1:GBP | |
|---|---|---|
| **Data collection and processing** | | |
| Magnification | 105,000 | |
| Voltage (kV) | 300 | |
| Electron dose (e-/Å$^2$) | 46 | |
| Defocus range (μm) | -0.9~-1.7 | |
| Pixel size (Å) | 0.835 | |
| Symmetry | C1 | |
| Initial particle images (no.) | 9,703,873 | |
| | $Ca_V1.2(\Delta C)/Ca_V\beta_3/Ca_V\alpha_2\delta$-1:GBP **(PDB:8FD7;EMD-29004)** | EMC:$Ca_V1.2(\Delta C)/Ca_V\beta_3$ **(EMD-29006**) |
| Final particle images (no.) | 259,107 | 383,185 |
| Map resolution (Å) | 3.1 | 3.1 |
| FSC threshold | 0.143 | 0.143 |
| Map resolution range (Å) | 2.4~8.0 | 2.4~8.0 |
| **Refinement** | | |
| Initial model used (PDB code) | 8EOG | |
| Model resolution (Å) | 3.3 | |
| FSC threshold | 0.5 | |
| Map sharpening *B* factor (Å$^2$) | -20 | |
| Model composition | | |
| Non-hydrogen atoms | 19,760 | |
| Protein residues | 2,416 | |
| Ligands | 24 | |
| *B* factors (Å$^2$) | | |
| Protein | 118.64 | |
| Ligand | 96.85 | |
| R.m.s deviations | | |
| Bond lengths (Å) | 0.008 | |
| Bond angles (°) | 1.171 | |
| Validation | | |
| MolProbity score | 2.08 | |
| Clashscore | 13.29 | |
| Poor rotamers (%) | 0.51 | |
| Ramachandran plot | | |
| Favored (%) | 93.11 | |
| Allowed (%) | 6.73 | |
| Disallowed (%) | 0.17 | |
